# Target cell adhesion limits macrophage phagocytosis and promotes trogocytosis

**DOI:** 10.1101/2025.02.06.636906

**Authors:** Kirstin Rollins, Sareen Fiaz, Meghan Morrissey

## Abstract

Macrophage phagocytosis is an essential immune response that eliminates pathogens, antibody-opsonized cancer cells and debris. Macrophages can also trogocytose, or nibble, targets. Trogocytosis and phagocytosis are often activated by the same signal, including IgG antibodies. What makes a macrophage trogocytose instead of phagocytose is not clear. Using both CD47 antibodies and a Her2 Chimeric Antigen Receptor (CAR) to induce phagocytosis, we found that macrophages preferentially trogocytose adherent target cells instead of phagocytose in both 2D cell monolayers and 3D cancer spheroid models. Disrupting target cell integrin using an RGD peptide or through CRISPR-Cas9 knockout of the αV integrin subunit in target cells increased macrophage phagocytosis. Conversely, increasing cell adhesion by ectopically expressing E-Cadherin in Raji B cell targets reduced phagocytosis. Finally, we examined phagocytosis of mitotic cells, a naturally occurring example of cells with reduced adhesion. Arresting target cells in mitosis significantly increased phagocytosis. Together, our data show that target cell adhesion limits phagocytosis and promotes trogocytosis.

## Introduction

Macrophages are innate immune cells that surveille the body for signs of injury or infection. Macrophages clear diverse targets, including pathogens, dying cells and debris, through phagocytosis (Freeman & Grinstein, 2021). Phagocytosis is critical for maintaining tissue homeostasis, protecting against infection and preventing autoimmunity (Gordon, 2016). Macrophages are also key effectors of many cancer therapies (Guerriero, 2018; Van Wagoner et al., 2023; Weiskopf & Weissman, 2015). Therapeutic antibodies like rituximab (CD20 antibody) and traztuzumab (Her2 antibody) bind to cancer cells and signal for macrophage uptake (Chao et al., 2010; Gül et al., 2014; Manches et al., 2003; Shi et al., 2015; Uchida et al., 2004; Weiskopf & Weissman, 2015). IgG is recognized by the Fc Receptor in macrophages (Nimmerjahn & Ravetch, 2008). Even many antibodies originally designed to block the function of their target benefit from activating the Fc Receptor (Chen et al., 2019; Dahan et al., 2016). More recently, engineering macrophages to express synthetic Chimeric Antigen Receptors (CARs) that trigger phagocytosis has been shown to shrink tumors in mouse models (Klichinsky et al., 2020; Morrissey et al., 2018; Sloas et al., 2021).

Activating phagocytosis of cancer cells is particularly exciting in solid tumors (Sloas et al., 2021). Macrophages are one of the most prevalent immune infiltrates of the solid tumor microenvironment (de Visser & Joyce, 2023). However, most studies on the mechanism of phagocytosis use suspended targets, including apoptotic cells, blood cancer cell lines, and reconstituted particles (Arandjelovic & Ravichandran, 2015; Gül et al., 2014; Joffe et al., 2020; Morrissey et al., 2018). Unlike these models, solid tumors have complex cell-cell and cell-ECM interactions that adhere cancer cells to the surrounding environment. How macrophages phagocytose a cell incorporated into a 3D tissue is not clear.

Phagocytosis of adherent targets often requires additional steps to remove the target from its environment. Adherent bacteria are pried from their substrate by a macrophage “hook-and-shovel” mechanism, wherein macrophages wedge lamellipodia between the bacteria and substrate to destroy adhesion molecules (Möller et al., 2013). Dying cells are loosened from their neighbors as focal adhesion and cell-cell contacts are dismantled during apoptosis (Brancolini et al., 1997). In melanoma models, macrophages cluster to cooperatively remove cancer cells (Dooling et al., 2023, 2024). These studies suggest that cell-cell and cell-substrate adhesion is a barrier to phagocytosis.

In addition to phagocytosis, macrophages can trogocytose, or nibble, target cells (Bettadapur et al., 2020). Trogocytosis has been observed in diverse contexts, including immune cell communication, developmental remodeling, and parasitic attack (Abdu et al., 2016; Gao et al., 2024; Hudrisier et al., 2001; Joly & Hudrisier, 2003; Mercer et al., 2018; Ralston et al., 2014; Weinhard et al., 2018). Trogocytosis and phagocytosis are triggered by the same signals, including IgG antibodies, and share many molecular regulators (Bettadapur et al., 2020; Lindorfer & Taylor, 2022). What makes a macrophage trogocytose instead of phagocytose is not clear.

Macrophages also trogocytose antibody opsonized cancer cells during cancer therapy (Beum et al., 2011; Lindorfer & Taylor, 2022; Park et al., 2022; Velmurugan et al., 2016). Trogocytosis of antibody opsonized cancer cells can lead to cancer cell death (Finotti et al., 2023; Matlung et al., 2018; Velmurugan et al., 2016). However, trogocytosis is also associated with ‘antigen shaving’, or the removal of target antigens, which limits antibody efficacy by making cancer cells more difficult for the immune system to detect (Beum et al., 2011; Hamieh et al., 2019; Kennedy et al., 2004; Williams et al., 2006). Biasing therapies towards phagocytosis could potentially improve cancer cell clearance (Williams et al., 2006; Zent et al., 2014).

In this study, we investigated how target cell adhesion impacts macrophage phagocytosis. We used an ovarian cancer cell line in 2D monolayers or 3D spheroids as a phagocytic target. Macrophages preferentially trogocytosed (nibbled) adherent cells and phagocytosed suspended cells. We show that this preference for trogocytosis was reversed if we reduced or eliminated cell-substrate adhesion. We also show that suspended cells were primarily trogocytosed instead of phagocytosed if we increased cell-cell adhesion. Finally, we observed that mitotic cells were more susceptible to phagocytosis than cells in interphase, indicating that macrophages may capitalize on opportune moments for phagocytosis upon the disassembly of adhesion. Together our results demonstrate that the macrophages preferentially trogocytose adherent targets.

## Results

### Construction of a Her2-targeting Chimeric Antigen Receptor

To study how macrophages attack adherent cells, we selected the SKOV3 ovarian cancer cell line as our target cell. This cell line has high expression of the human epidermal growth hormone receptor 2 (Her2/ErbB2). Traztusubmab, an antibody targeting Her2, is a common cancer therapy and has previously been shown to induce cancer cell internalization (Petricevic et al., 2013; Shi et al., 2015; Velmurugan et al., 2016). Her2-targeting CAR macrophages are being investigated in clinical trials (Klichinsky et al., 2020; Sloas et al., 2021).

We elected to induce phagocytosis with a Her2-targeted CAR, since Her2 antibodies inhibit Her2 signaling in addition to activating phagocytosis (Hudziak et al., 1989; Swain et al., 2023). We designed a Her2-targeting CAR comprised of an extracellular Her2 antibody fragment (scFv from 4D5-8 (Carter et al., 1992)), the CD8 hinge and transmembrane domain from successful CAR T molecules (Fesnak et al., 2016; Kochenderfer et al., 2009), the intracellular signaling domain from the mouse Fc Receptor common gamma chain (Kern et al., 2021; Morrissey et al., 2018), and a GFP tag (Fig. 1A). A similar Her2 CAR was recently shown to decrease cancer growth in mouse xenograft models lacking T cells, suggesting that Her2 CAR macrophages can shrink tumors without engaging the adaptive immune system (Klichinsky et al., 2020).

**Figure 1:**
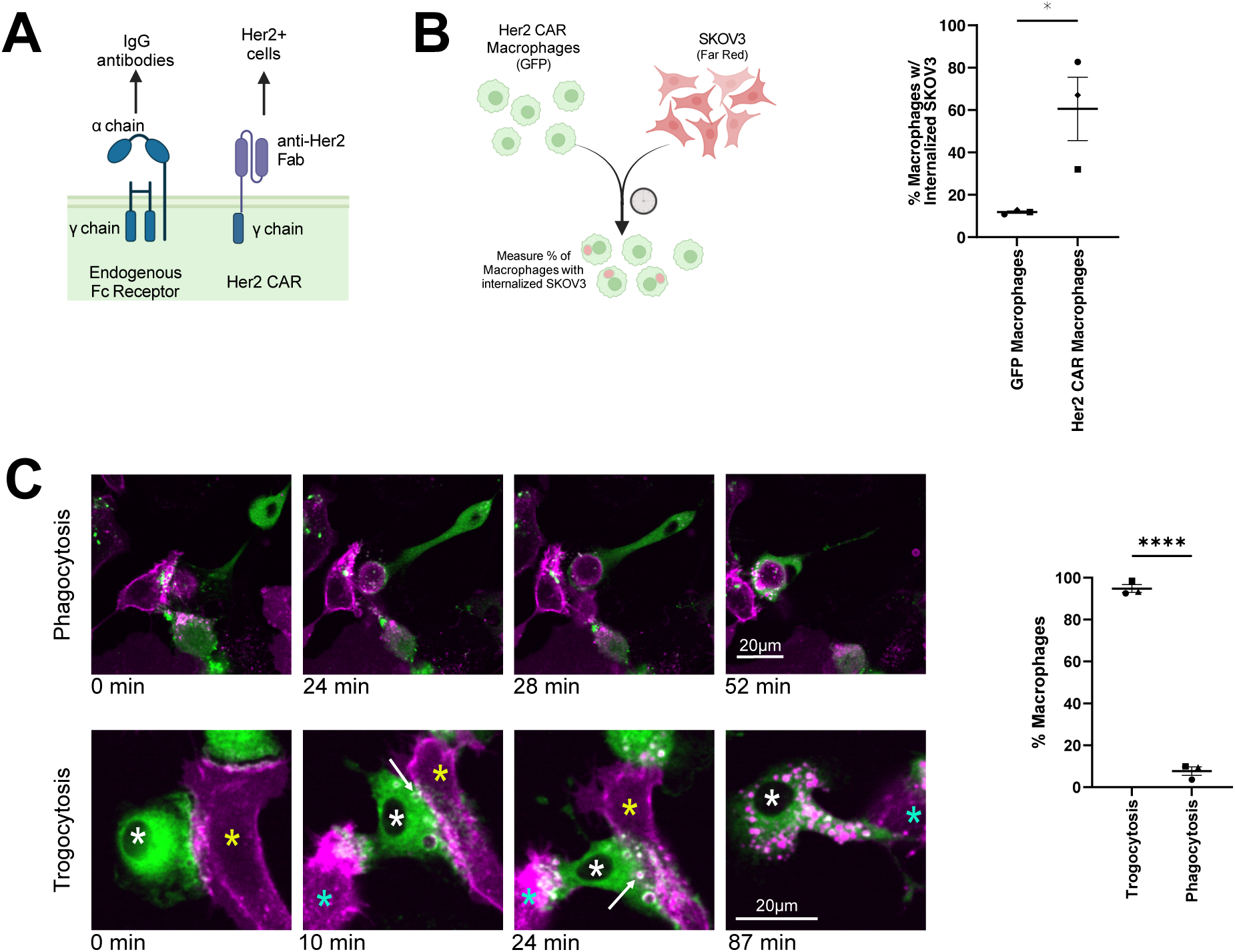
Her2 CAR macrophages trogocytose target SKOV3 cells. (A) Schematic shows the design of the Her2 CAR (right) compared to the native Fc Receptor (left). The Her2 CAR contains an extracellular scFv recognizing Her2, and activates phagocytosis via the intracellular signaling domain from the endogenous Fc Receptor common gamma chain. (B) Schematic (left) describes the assay quantified in the graph (right). Her2 CAR GFP or control GFP (GFP-CAAX) bone marrow derived macrophages (BMDMs) were mixed with SKOV3 cells dyed with CellTrace Far Red. Flow cytometry was used to measure the percent of macrophages (GFP+) that internalized cancer cell material (Far Red+). (C) Her2 CAR GFP (green) macrophages were visualized interacting with SKOV3 cancer cells (mCh-CAAX; magenta) using timelapse confocal microscopy. Stills from the images are depicted on the left, and the percent of macrophages engaging in trogocytosis (internalizing part of a cell) or phagocytosis (internalizing a whole cell) is graphed to the right. These stills correspond to Video 1 (top) and Video 2 (bottom). In B and C, data was compared using a Students T test. N= 3 experiments with independently generated and infected BMDMs. In all graphs, bars represent the mean ± SEM. Data collected on the same day is annotated with the same shape point. * denotes p<0.05, **** denotes p<0.0005. Scale bar denotes 20 μm.

To test whether the Her2 CAR could stimulate macrophage activity, we incubated Her2 CAR GFP or control GFP macrophages with Her2+ SKOV3 ovarian cancer cells dyed in Far Red Cell Trace. After 2 hours, we measured the cancer cell uptake via flow cytometry. We found that Her2 CAR expression increased macrophage internalization of SKOV3 cells (Fig. 1B).

### Her2 CAR macrophages primarily trogocytose target cells

To further characterize macrophage activity against adherent targets, we used confocal live-cell timelapse imaging of Her2 CAR macrophage and SKOV3 co-incubations. SKOV3 cells were infected with a membrane tethered mCherry (mCherry-CAAX). To our surprise, phagocytosis of cancer cells was relatively rare. We observed that only 10% of macrophages phagocytosed during our 10 hour timelapse (Fig. 1C, Video 1). In contrast, 95% of macrophages trogocytosed, or nibbled, the cancer cell membrane (Fig. 1C, Video 2). This data shows that most SKOV3 internalization is trogocytosis, rather than phagocytosis.

### Her2 CAR macrophages limit the growth of SKOV3 spheroids

Our previous experiments in monolayers lacked the 3D architecture and cell-to-cell adhesion normally found in a solid tumor. To investigate macrophage eating dynamics in a simple 3D tissue model, we built spheroids of SKOV3 cells. To generate spheroids, we plated membrane labeled (mCh-CAAX) SKOV3 cells in a low adhesion dish in media supplemented with 2.5% matrigel to favor cell-cell interactions (Ivascu & Kubbies, 2007; Heredia-Soto et al., 2018; Tofani et al., 2020). By 72 hours, the SKOV3 cells assembled into consistent, compact spheroids (445 ± 65 μm in diameter). We then transferred the pre-assembled SKOV3 spheroids into a matrigel dome containing Her2 CAR or control GFP macrophages. Z-stack images showed that both GFP and Her2 CAR macrophage could invade into the spheroids within 24 hours (Videos 3,4). We tracked spheroid size for 10 days via spinning disc confocal imaging (Fig. 2A,B). We found that Her2 CAR macrophages significantly reduced the size of the spheroids compared to no macrophages or GFP macrophages (Fig. 2B). In the final timepoints, a large portion of the remaining mCherry signal appeared to be from macrophages that had internalized cancer cells (Fig. 2C). This demonstrates that Her2 CAR macrophages effectively attack 3D cancer spheroids.

**Figure 2:**
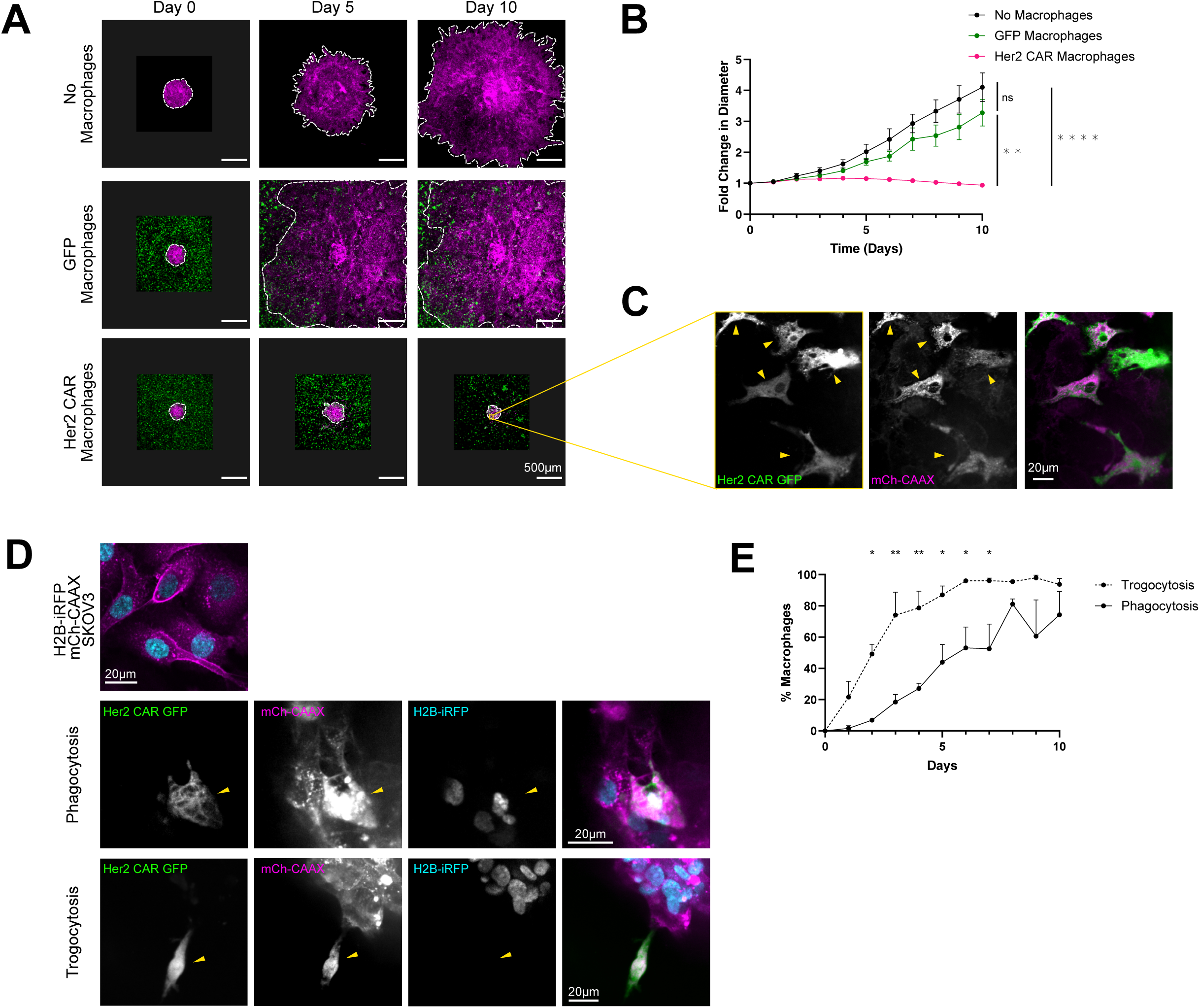
Her2 CAR macrophages trogocytose more than phagocytose in 3D spheroid models. (A) SKOV3 cells (mCh-CAAX, magenta) were assembled into a 3D spheroid and embedded in matrigel alone (top), with control GFP (green) macrophages (middle) or with Her2 CAR GFP (green) macrophages (bottom). Images are maximum projections of spinning disc confocal z stacks. Dashed line highlights the spheroid boundary. Scale bars denote 500 μm. On days 5 and 10, multiple images were stitched together to capture the entire spheroid in the no macrophage and GFP macrophage conditions. Unstitched images are presented on a grey background so the image scale is consistent. (B) Graph depicts spheroid growth over time for the same conditions shown in (A). Individual spheroids were tracked for 10 days, and the diameter was normalized to the starting diameter. The length and width of the spheroid were measured on the max projection of confocal z-stacks and averaged to calculate spheroid diameter. N=6 spheroids acquired in 4 independent experiments. (C) Inset from (A) shows that mCherry signal on day 10 is mostly contained within Her2 CAR GFP (green) macrophages. Arrowheads point to macrophages. Scale bars denote 20 μm. (D) Top image shows the H2B-iRFP (blue), mCherry-CAAX (magenta) SKOV3 cell line used to assemble spheroids. Images show examples of Her2 CAR GFP (green) macrophages that have phagocytosed (middle, internalized nuclei position is highlighted with yellow arrowhead) or trogocytosed (bottom, yellow arrowhead) SKOV3 cells. Scale bars denote 20 μm. (E) The percent of Her2 CAR GFP macrophages that had phagocytosed (internalized H2B-iRFP) or trogocytosed (internalized mCh-CAAX) was quantified on each day. Only macrophages touching the spheroid were included in the analysis. N= 3 spheroids from 3 independent experiments. In B, data acquired on day 10 was compared with a one-way ANOVA with Holm-Sidak’s multiple comparison test. In E, data from each day was compared with a two-way ANOVA. Bars represent the mean ± SEM * denotes p<0.05, ** denotes p<0.005, **** denotes p<0.00005.

### Macrophages trogocytose more than phagocytose in a 3D environment

We next sought to determine if macrophages attack SKOV3 spheroids via trogocytosis as in the 2D culture system. Brief timelapse imaging revealed several examples of trogocytosis and phagocytosis but was difficult to quantify (Video 5, Fig. S1). Instead, we double-labeled cancer cells with a membrane tethered mCherry (mCh-CAAX) and a nuclear iRFP (H2B-iRFP; Fig. 2D). If a cancer cell were phagocytosed, both nuclear and membrane signals would be detected within the macrophage, confirming whole cell engulfment. If a cancer cell were trogocytosed, only the membrane signal would be detected within the macrophage. We imaged the spheroids daily for 10 days and quantified the fraction of macrophages that had internalized mCherry (trogocytosed) or iRFP and mCherry (phagocytosed) (Fig. 2D,E). We found that trogocytosis was more common than phagocytosis in 3D spheroid models (Fig. 2E), as in the 2D culture system. Trogocytic macrophages were clearly observed in the first days after macrophage infiltration, while phagocytic macrophages slowly accumulated over time (Fig. 2E).

### Macrophage trogocytosis strips the antigen from target cells

Several clinical studies have noted that trogocytosis of antibody-opsonized cancer cells removes target antigen from the cancer cell surface in vivo (Hamieh et al., 2019; Lindorfer & Taylor, 2022; Park et al., 2022; Williams et al., 2006). This process, known as antigen shaving, makes the cancer cells more difficult for immune cells to detect (Hamieh et al., 2019; Lindorfer & Taylor, 2022; Williams et al., 2006). We checked for antigen shaving by immunostaining for Her2 on SKOV3 cells after macrophage treatment. We found that treatment with Her2 CAR macrophages led to a substantial decrease in the surface levels of Her2, whereas GFP macrophages did not change Her2 surface levels (Fig. 3A,B). Overall, this data illustrates the high level of trogocytosis in our system.

**Figure 3:**
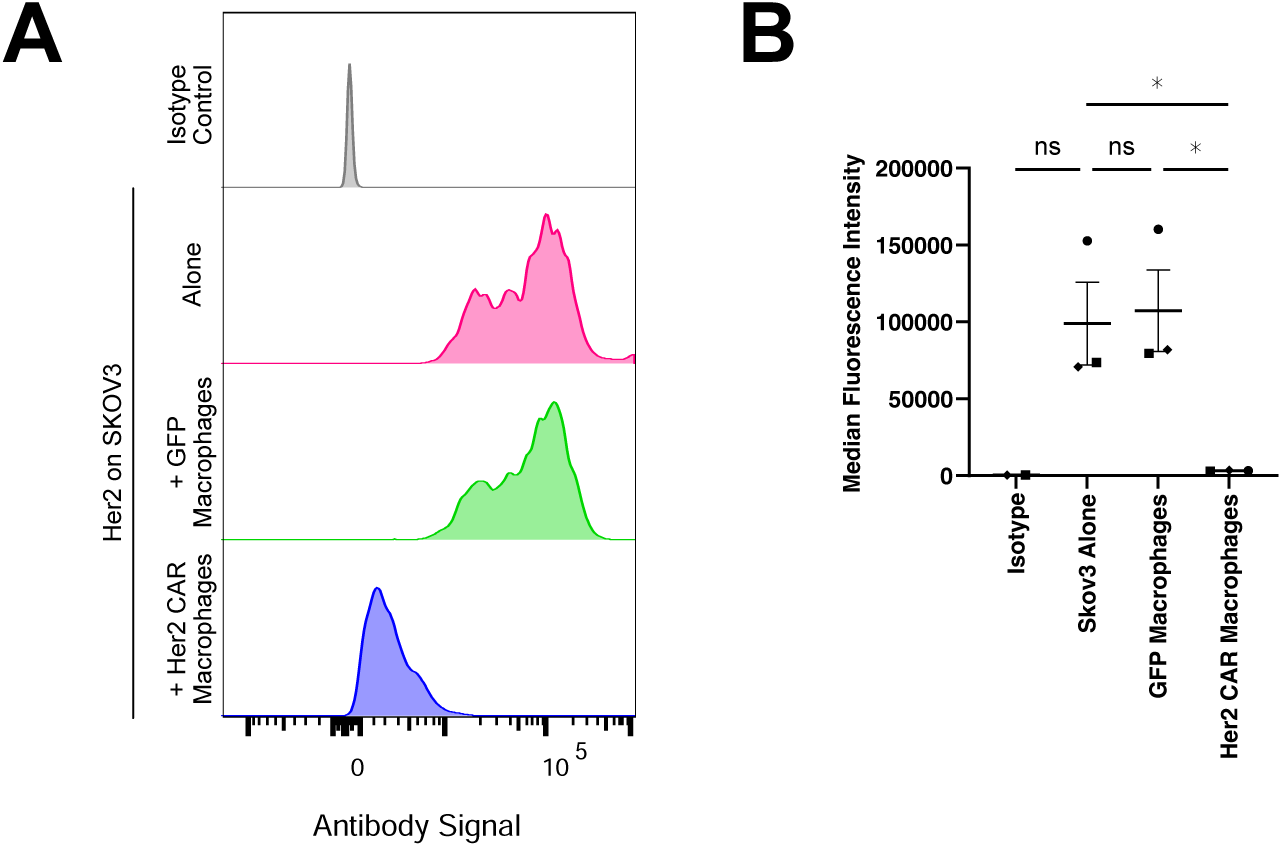
Trogocytosis strip Her2 from SKOV3 target cells. (A) Her2 CAR GFP or control GFP macrophages were incubated with SKOV3 cells (mCh-CAAX) for 2 hours. The cells were then stained for Her2 and analyzed by flow cytometry. A representative flow plot shows the Her2 levels on SKOV3 cells alone (pink), and SKOV3 cells incubated with Her2 CAR (blue) or control GFP macrophages (green). SKOV3 cells were also stained with an isotype control antibody as a negative control (grey). (B) Graph shows the median fluorescence intensity from three independent replicates. Bars represent the mean ± SEM of the replicates. Data was compared using a one way ANOVA with a Holm-Sidak multiple comparison test. Data collected on the same day is annotated with the same shape point. * denotes p<0.05.

### Macrophages primarily phagocytose suspended cells and trogocytose adherent cells

Prior studies using antibody opsonized B cells have reported high levels of phagocytosis (for example, (Chao et al., 2010; Gül et al., 2014)). We have previously seen that CD19 CAR macrophages targeting B cells phagocytose many cancer cells during a similar timelapse microscopy experiment (Mishra et al., 2023; Morrissey et al., 2018). We hypothesized that adherent cells may be more difficult to phagocytose and more likely to be trogocytosed than suspended cells. To see if this was broadly true for multiple adherent cell lines, we assembled a panel of cancer cell lines, including three adherent cell lines and two suspension cell lines. We expressed membrane tethered mCherry-CAAX and a nuclear H2B-iRFP marker in all the cell lines, opsonized the cells with a CD47 antibody, and measured trogocytosis and phagocytosis with flow cytometry (Fig. 4A). The selected cancer cell lines expressed a reasonably uniform level of CD47, a ‘Don’t Eat Me’ signal (Fig. S2). CD47 antibodies trigger phagocytosis by blocking the inhibitory CD47 signal and engaging with macrophage Fc Receptors through the antibody Fc domain (Chao et al., 2010; Jaiswal et al., 2009; Majeti et al., 2009; Osorio et al., 2023). We found that the total number of macrophages internalizing any cancer cell material (mCherry positive) was relatively uniform across the five cell lines (Fig. 4B). Strikingly, we observed that the suspended cell lines were primarily phagocytosed (60-70% of total activity) and the adherent cell lines were primarily trogocytosed (80-90% of total activity; Fig. 4C). Since antibody opsonization induced trogocytosis of multiple adherent cell lines, this demonstrates that trogocytosis is not a unique feature of the SKOV3 target cell, or of the CAR macrophage system.

**Figure 4:**
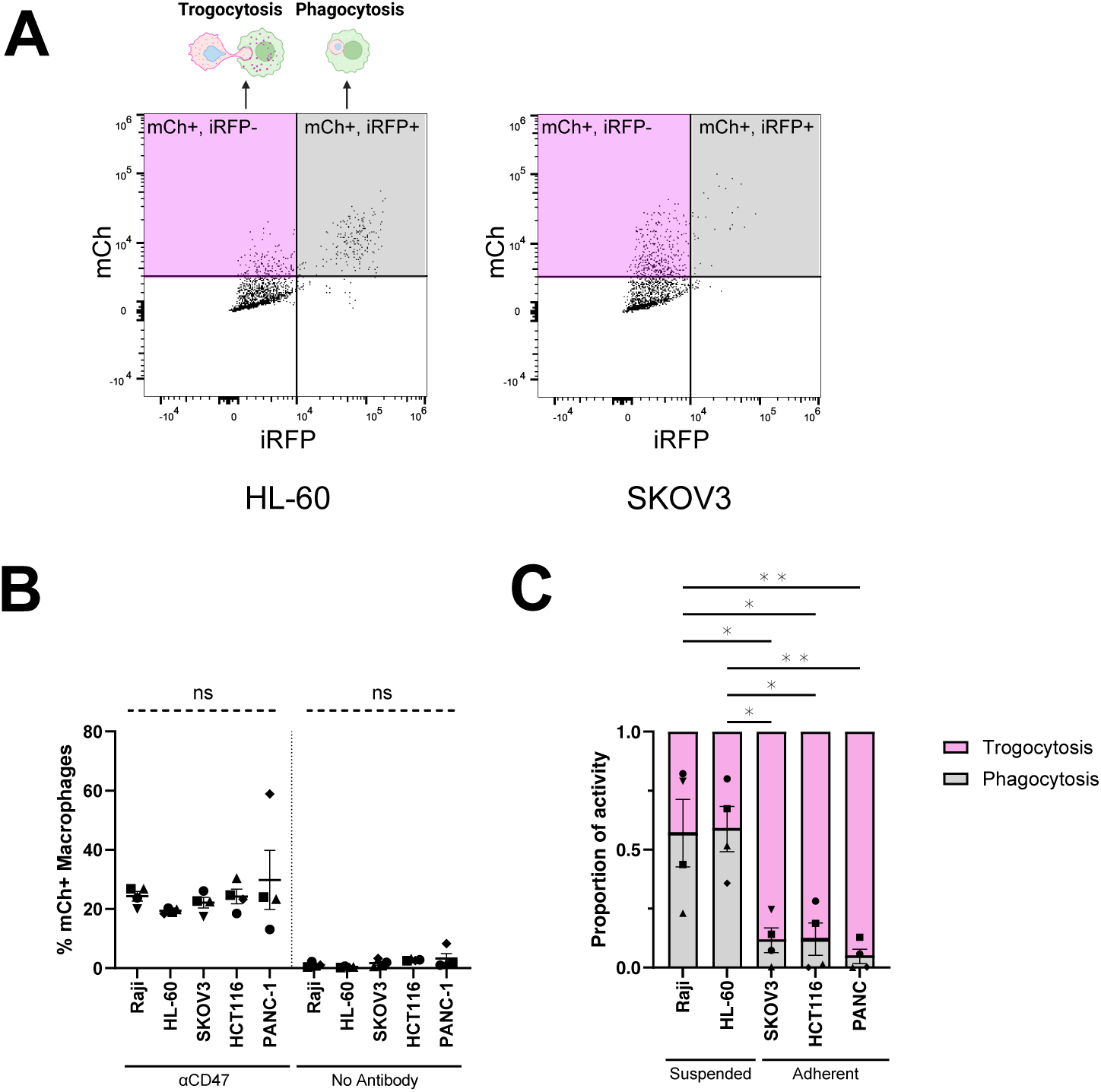
Trogocytosis is more common than phagocytosis in multiple adherent cancer cell lines. Cancer cells were opsonized with a mouse anti-human CD47 antibody, and the amount of phagocytosis and trogocytosis was measured by flow cytometry. (A) Representative flow plots show the assay for distinguishing phagocytosis and trogocytosis by flow cytometry. (B) Graph shows the percent of mCherry positive macrophages, indicating that these macrophages internalized some cancer cell material. (C) Graph shows the fraction of mCherry positive macrophages that were also positive for iRFP, indicating cancer cell phagocytosis (grey), or lacking iRFP, indicating trogocytosis (pink). For B and C, data was compared using one-way ANOVA with Holm-Sidak multiple comparison correction. N = 4 independent experiments, consisting of 3 averaged technical replicates. Bars represent the mean ± SEM. Data collected on the same day is annotated with the same shape point. * denotes p<0.05, ** denotes p<0.005.

### Decreasing target cell adhesion increases phagocytosis

We hypothesized that decreasing target cell adhesion could shift macrophage behavior from trogocytosis to phagocytosis. First, we tested if detaching SKOV3 cells from a substrate increased phagocytosis. We compared phagocytosis of adherent SKOV3 cells to phagocytosis of detached SKOV3 cells. Using timelapse microscopy to measure phagocytosis, we found Her2 CAR macrophages phagocytosed more SKOV3 cells when the cells were presented in suspension than when allowed to adhere to a tissue culture plate (Fig. 5A, Fig. S3). This was true whether the macrophages were mixed with the SKOV3 cells in suspension, or pre-adhered to the tissue culture plate (Fig. S3). Control GFP macrophages did not phagocytose more suspended cells, suggesting this increase in phagocytosis was due to Her2 CAR activity not increased cell death (Fig. 5A). Together, these data show that suspended cells are more readily phagocytosed than adherent cells.

**Figure 5:**
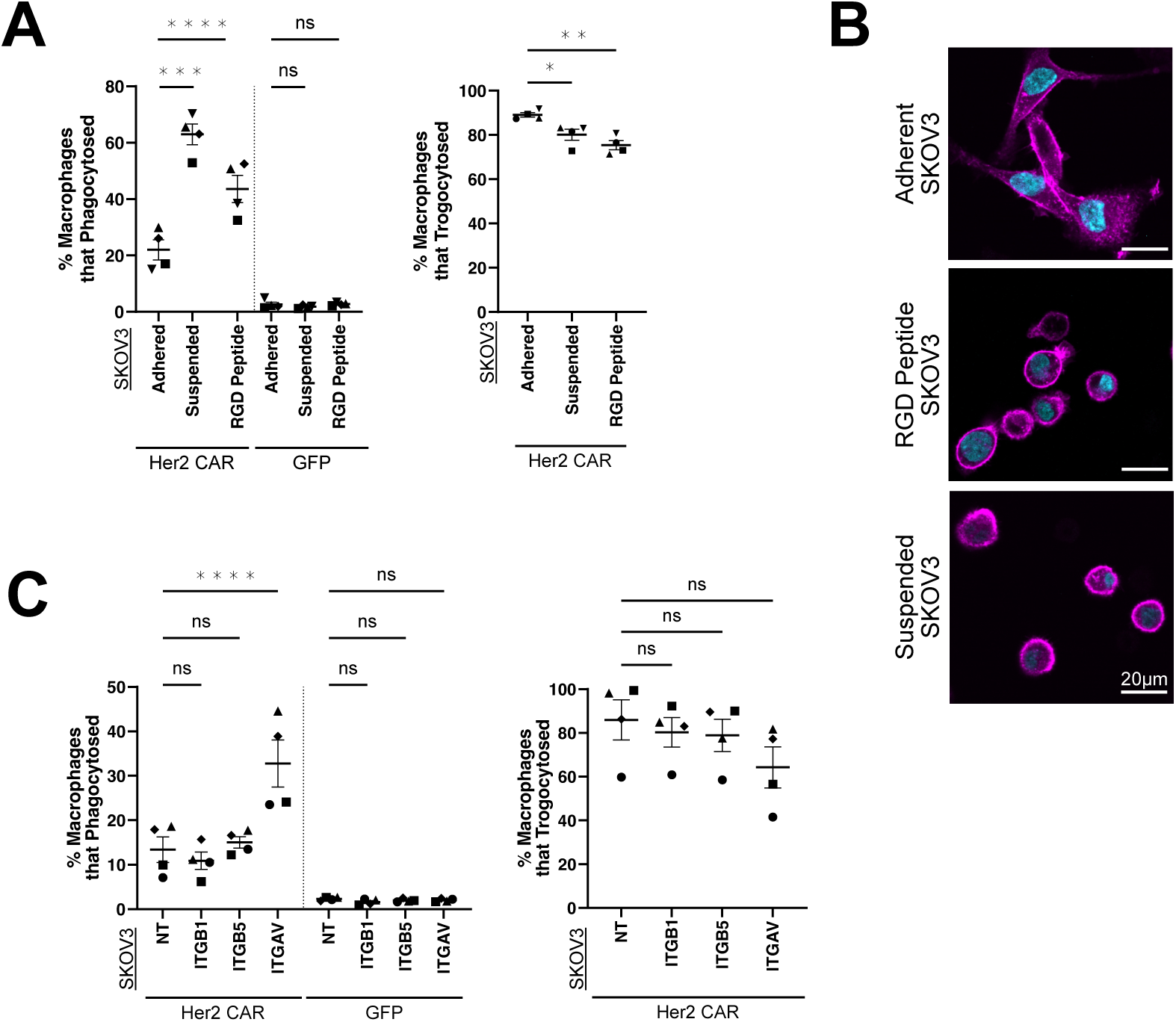
Reducing cancer cell adhesion promotes phagocytosis. (A) Her2 CAR GFP or control GFP-CAAX macrophages were incubated with adherent SKOV3 cells, RGD peptide treated SKOV3 cells, or suspended SKOV3 cells for 8 hours. The percent of macrophages that phagocytosed (right) and trogocytosed (left) was measured during an 8 hour timelapse. (B) SKOV3 cells with a membrane tethered mCherry (mCh-CAAX) and a nuclear iRFP (H2B-iRFP) were plated with 50 μg/mL RGD peptide 8 hours prior to macrophage addition to block adhesion. (C) SKOV3 cells expressing mCherry-CAAX and H2B-iRFP were infected with CRISPR-Cas9 and a sgRNA targeting integrin β1, β5, αV or a non targeting control guide (NT). Infected cells were selected with puromycin and knockdown was confirmed by flow cytometry (Fig. S4). The resulting polyclonal cell lines were incubated with Her2 CAR GFP or control GFP-CAAX macrophages and the percent of macrophages that phagocytosed (left) and trogocytosed (right) was measured during an 8 hour timelapse. Data was compared using one-way ANOVA with Holm-Sidak multiple comparison correction. N = 4 independent experiments. Bars represent the mean ± SEM. Data collected on the same day is annotated with the same shape point. * denotes p<0.05, ** denotes p<0.005, *** denotes p<0.0005, **** denotes p<0.00005. Scale bars denote 20 μm.

We next sought to modulate cell-substrate attachment by decreasing integrin adhesion. To weaken cell-substrate adhesion, we treated SKOV3 cells expressing membrane tethered mCherry and nuclear iRFP with the RGD peptide, Cilengitide Trifluoroacetate. This RGD peptide primarily inhibits the integrins a_V_β_3_ and a_v_β_5_ (Dechantsreiter et al., 1999). We verified that the RGD peptide caused the SKOV3 cells to adopt a rounded morphology and decreased surface area characteristic of lowered adhesion, although the cells were not completely suspended (Fig. 5B). To minimize the RGD peptide’s effect on the macrophages, we washed out the RGD peptide before adding Her2 CAR or control GFP macrophages. We then measured phagocytosis and trogocytosis by live-cell timelapse microscopy. We found that RGD peptide promoted phagocytosis and reduced trogocytosis, although fewer cells were phagocytosed than when the SKOV3 cells were fully detached from the substrate (Fig. 5A). Again, control GFP macrophages did not phagocytose additional target cells in the RGD condition, suggesting that these cells are not exposing additional pro-phagocytic signals. This suggests that inhibiting integrin is sufficient to increase phagocytosis.

To confirm that inhibiting integrin in the target cell increases phagocytosis, we used CRISPR-Cas9 to knock out specific integrin subunits in SKOV3 cells (Fig. S4, Table S1). We selected integrin β1 and β5 (ITGB1, ITGB5) as dysregulation of these receptors is associated with cancer development and progression in a wide variety of solid tumors, including Her2+ breast and ovarian cancers (Casey et al., 2001; Davidson et al., 2003; Desgrosellier & Cheresh, 2010). We also selected integrin αV (ITGAV) as it is reported to affect ovarian cancer proliferation and metastasis (Casey et al., 2001; Cruet-Hennequart et al., 2003; Davidson et al., 2003). ITGAV knockout target cells were phagocytosed more than control cells (Fig. 5C). ITGB1 and ITGB5 knockouts did not significantly impact phagocytosis, which could indicate redundancy since ITGAV pairs with both B subunits. These findings demonstrate decreasing cell-substrate adhesion promotes phagocytosis.

### Increasing cell-cell adhesion in suspension cells decreases phagocytosis

We next wondered if we could conversely increase adhesion to limit phagocytosis. We overexpressed E-Cadherin (CDH1) in Raji B cells, a suspension cell line. Raji B cells expressing E-Cadherin formed multicellular clusters, indicating increased cell-cell adhesion (Fig. 6A). We opsonized wild type and E cadherin overexpressing Rajis with a mouse anti-human CD47 antibody to induce phagocytosis. Macrophages with Rajis expressing either iRFP-CAAX or H2B-iRFP were co-incubated for 2 hours before dissociating the clustered conditions into a single-cell solution for flow cytometry. The level of phagocytosis (H2B-iRFP internalization) was significantly lower in the E-Cadherin overexpressing clusters than the wild type Raji cells (Fig. 6B). In contrast, the fraction of macrophages with internalized Raji cell membrane (iRFP-CAAX), either by trogocytosis or phagocytosis, did not change (Fig. 6C). This suggests that macrophages encountered a similar number of Raji targets, but did not phagocytose the E cadherin overexpressing cells. Overall, this demonstrates that increasing cell-cell adhesion in target cells limits phagocytosis.

**Figure 6:**
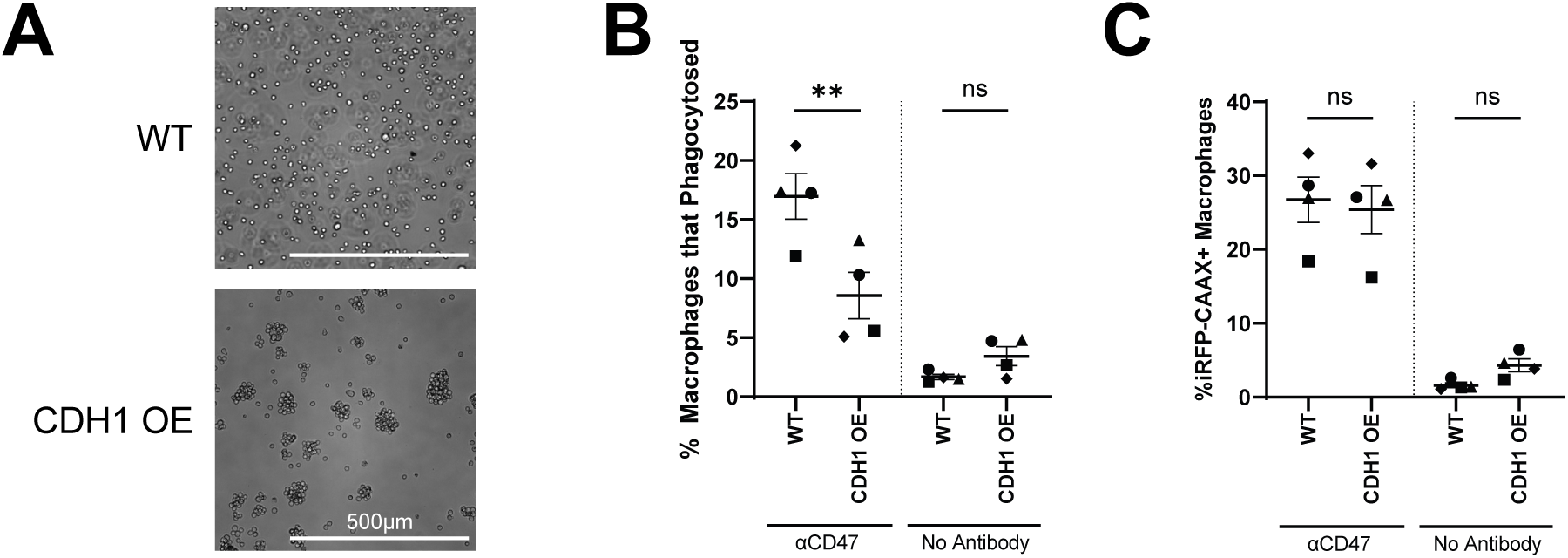
Increasing cancer cell adhesion reduces cancer cell phagocytosis. (A) Raji B cells were infected with E cadherin GFP (CDH1 OE) to induce cell-cell adhesion. Bright field images show the formation of cell clusters in CDH1 overexpressing cells. (B) Raji B cells expressing H2B-iRFP were opsonized with a CD47 antibody and incubated with mCh-CAAX macrophages for 2 hours. The percent of phagocytic macrophages was measured by flow cytometry. Scale bars denote 500 μm. (C) Raji B cells expressing iRFP-CAAX were opsonized with a CD47 antibody and incubated with mCh-CAAX macrophages for 2 hours. The percent of iRFP positive macrophages was measured by flow cytometry. Data was compared using one-way ANOVA with Holm-Sidak multiple comparison correction. N = 4 independent experiments. Bars represent the mean ± SEM. Data collected on the same day is annotated with the same shape point. ** denotes p<0.005.

### Dividing cells are more vulnerable to phagocytic attack

Our data demonstrates that macrophages phagocytose cellular targets with weakened attachment to substrates and neighboring cells. However, even without any synthetic manipulations of target cell adhesion, we observed that 10% of macrophages could phagocytose adherent SKOV3 cells (Fig. 1E). We revisited these intrinsic phagocytic attacks in our timelapses and found that a large number of the phagocytosed cells appeared to be in the process of cell division (Fig. 7A, Video 6). While only 4% of SKOV3 cells were dividing at any time, 38% of the phagocytosed SKOV3 cells were undergoing division (Fig. 7B). Mitotic cells are known to have reduced adhesion (Akhmanova et al., 2022; Dix et al., 2018; Lock et al., 2018; Marchesi et al., 2014), so this observation is consistent with our overall hypothesis that reduced cell adhesion promotes phagocytosis.

**Figure 7:**
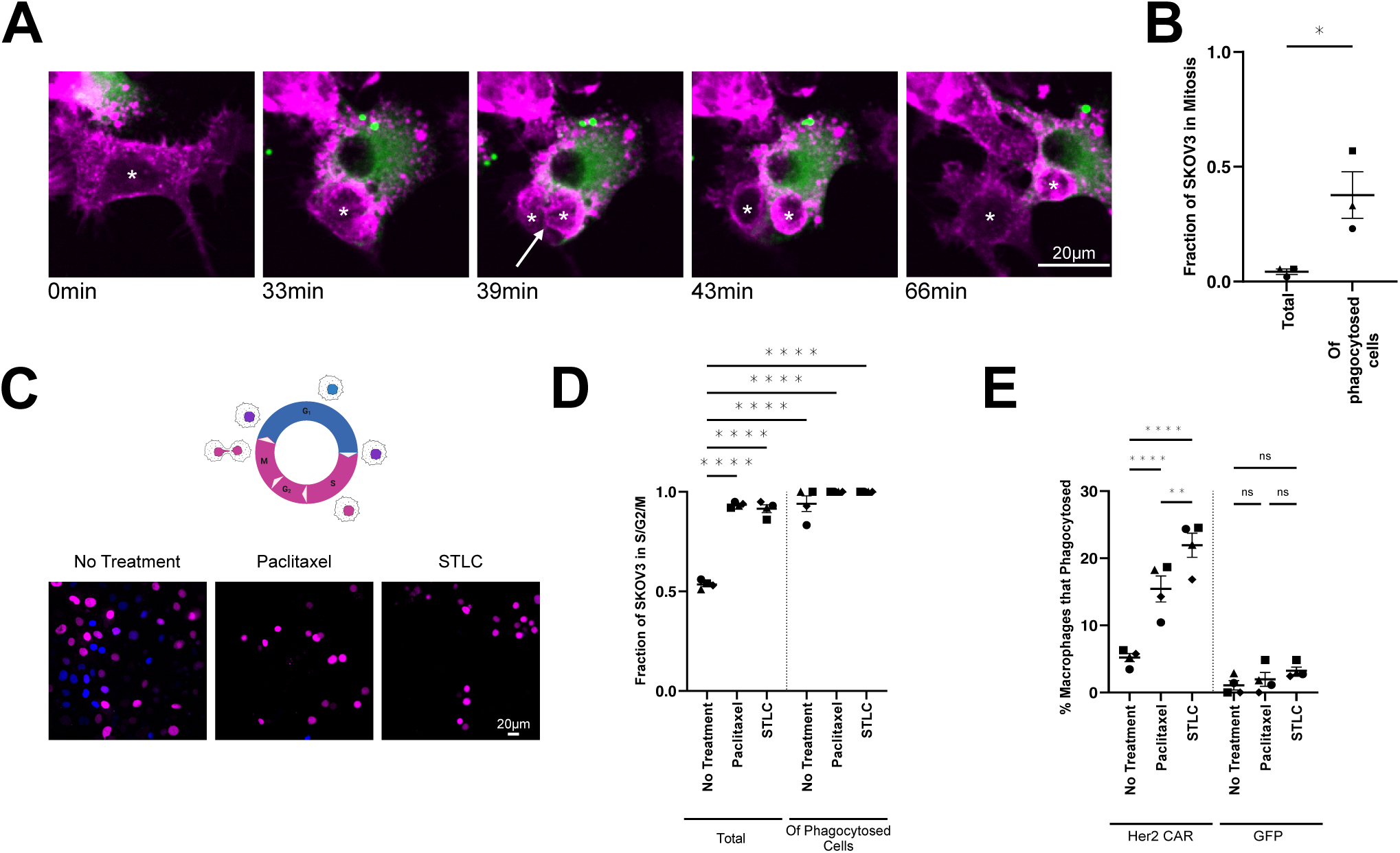
Mitotic cells are phagocytosed more than their interphase neighbors. (A) Stills from Video 6 show a Her2 CAR GFP (green) macrophage phagocytosing a mitotic SKOV3 (mCherry-CAAX, magenta). (B) Graph shows the fraction of SKOV3 undergoing division in the entire population (left) compared to the fraction of phagocytosed SKOV3 cells undergoing division (right). (C) SKOV3 cells expressing FUCCI were treated with 0.1 μM paclitaxel or 1 μM STLC for 20 hours to arrest cells in mitosis. Image shows cells in G1 phase (blue; hCdt1) and S/G2/M (magenta; hGeminin). (D) SKOV3 cells were treated with paclitaxel or STLC for 20 hours, then the inhibitors were removed and replaced with fresh media and Her2 CAR GFP macrophages. The fraction of mCherry+ FUCCI SKOV3 cells (indicating S/G2/M phase) was measured for the entire SKOV3 population (Total) and for the SKOV3 cells that were phagocytosed (Of Phagocytosed Cells) during a 4 hour timelapse. (E) Graph shows the percent of Her2 CAR GFP or GFP-CAAX macrophages phagocytosing a cancer cell in the same experiment. For B, data was compared using a Students T test. N= 3 independent experiments. For D and E, data was compared using one-way ANOVA with Holm-Sidak multiple comparison correction. N = 4 independent experiments. Bars represent the mean ± SEM. Data collected on the same day is annotated with the same shape point. * denotes p<0.05, ** denotes p<0.005, **** denotes p<0.00005. Scale bars denote 20 μm.

To examine this more closely, we generated a SKOV3 cell line expressing Fluorescent Ubiquitination-based Cell Cycle Indicator (FUCCI) to read out the stage of the cell cycle. The FUCCI reporter consists of an mCherry tagged hGeminin that will accumulate in S/G2/M before being degraded, and a BFP tagged hCdt1 will be detected in G1 (Fig. 7C)(Sakaue-Sawano et al., 2008; Sato et al., 2019). Using this reporter, we found that 94% of phagocytosed SKOV3 were mCherry positive, indicating that these cells were in S/G2/M, while only 53% percent of the overall SKOV3 population was mCherry positive (Fig. 7D). This is consistent with our observation that SKOV3 cells are more likely to be phagocytosed while dividing.

We next hypothesized that arresting target cells in mitosis would increase phagocytosis. To test this, we treated cancer cells with paclitaxel (Taxol) or S-trityl-cysteine (STLC). Paclitaxel is a cancer chemotherapeutic that inhibits microtubule disassembly, and prevents cells from passing the spindle assembly checkpoint (Horwitz, 1994). STLC inhibits Eg5, a mitotic kinesin necessary for assembly of the bipolar spindle, locking the cells in prometaphase (Skoufias et al., 2006). After 20 hours of paclitaxel and STLC treatment, nearly all FUCCI SKOV3 cells were mCherry positive indicating cell cycle arrest (Fig. 7C,D). The mitotic arrest did not induce apoptosis by 20 hours, as we did not detect an increase in phosphatidylserine exposure (Fig. S5). We then co-incubated the arrested cells with Her2 CAR macrophages and measured phagocytosis by timelapse microscopy. We found that cancer cells arrested in mitosis were phagocytosed significantly more than untreated controls (Fig. 5E). This data suggests that mitotic cells are easier to phagocytose than interphase cells.

## Discussion

While many antibody therapies and CAR macrophages are presumed to primarily trigger phagocytosis, we have found that phagocytosis of adherent cells is relatively rare. Instead, macrophages trogocytose, or nibble, these cells. Reducing target cell adhesion by disrupting integrin increased macrophage phagocytosis and reduced trogocytosis. Conversely, increasing cell-cell adhesion in a suspended cell line reduced phagocytosis. Overall, this data shows that target cell adhesion regulates phagocytosis and trogocytosis.

Very little is known about what makes a macrophage trogocytose instead of phagocytose. Trogocytosis and phagocytosis are controlled by nearly identical signaling pathways in the macrophage (Bettadapur et al., 2020). Trogocytosis and phagocytosis also occur on diverse targets, making it unlikely that there is a specific molecule that signals for trogocytosis instead of phagocytosis (Bettadapur et al., 2020). Instead, the biophysical properties of the target cell may guide the macrophage to trogocytose instead of phagocytose. A recent study found that target membrane tension is a key regulator of trogocytosis (Cornell et al., 2024). Lower membrane tension increased trogocytosis, while higher membrane tension favored phagocytosis. Our study also supports the hypothesis that the target’s biophysical properties determine if a macrophage performs phagocytosis or trogocytosis.

Trogocytosis has been observed in many clinical settings (Hamieh et al., 2019; Lindorfer & Taylor, 2022; Park et al., 2022; Williams et al., 2006). The impact of trogocytosis is murky. Sometimes, trogocytosis kills the target cell (Finotti et al., 2023; Matlung et al., 2018; Park et al., 2022; Velmurugan et al., 2016). In other cases, trogocytosis limits the efficacy of cancer therapy. Trogocytosis of the CD20 antibody therapy rituximab causes a significant reduction in CD20 levels on cancer cells in rituximab-treated patients (Kennedy et al., 2004). Optimizing the rituximab dosing schedule to limit trogocytosis improved efficacy (Williams et al., 2006; Zent et al., 2014). Similarly, high affinity CAR T cells trogocytose target antigen from cancer cell’s surface leading to antigen escape (Hamieh et al., 2019). Re-engineering the CAR T molecule to have a lower affinity increased efficacy by reducing antigen shaving. These studies suggest that biasing antibody therapies towards phagocytosis instead of trogocytosis could significantly reduce antigen shaving and improve clinical efficacy.

We found that arresting cancer cells in mitosis increased CAR macrophage phagocytosis. Dividing cells disassemble focal adhesion, while remaining loosely attached to their underlying substrate (Dix et al., 2018). This lowered adhesion allows macrophages to squeeze past dividing cells and infiltrate cell-dense tissue (Akhmanova et al., 2022). Dividing cells undergo other cell shape changes that may promote phagocytosis, including cell rounding. However, given our data showing that decreasing cell adhesion promotes phagocytosis, we attribute at least part of the increased phagocytosis to decreased cell adhesion. Interestingly, a combination of paclitaxel and Her2 monoclonal antibody is a highly effective treatment for Her2 positive breast cancers (Giordano et al., 2022). This suggests that the current therapy regimen could already benefit from enhanced phagocytosis due to mitotic arrest.

We also demonstrate that reducing integrin-mediated adhesion in target cells increased macrophage phagocytosis. Integrin inhibitors have previously been targeted in several clinical trials (Desgrosellier & Cheresh, 2010). Unfortunately, many of these integrin inhibitors failed in clinical trials due to lack of efficacy. Our work suggests that while these inhibitors were ineffective as a monotherapy, they may synergize with antibody therapies that induce phagocytosis.

In some cases, trogocytosis is reported to kill cancer cells. Why trogocytosis is sometimes lethal and sometimes not is unclear. Several of the studies showing death after trogocytosis induce trogocytosis with a Her2 antibody (Matlung et al., 2018; Velmurugan et al., 2016). We find that trogocytosis by Her2 CAR macrophages dramatically decreases Her2. Our study measured Her2 levels on viable cancer cells, suggesting that this reduction in Her2 did not immediately kill the cancer cells. However, this reduction in Her2 likely decreases cancer cell fitness since Her2 provides a powerful pro-growth signal to Her2+ cancers. Future studies will need to address the impact of trogocytosis on cancer cell fitness.

Macrophages co-operate to phagocytose in solid tumor models (Dooling et al., 2023, 2024). Our studies support the idea that cell-cell and cell-matrix adhesions are too strong for a single macrophage to overcome during phagocytosis (Dooling et al., 2023). We speculate that trogocytosis could be a mechanism for macrophage cooperation by weakening target cells. Our data shows higher occurrence of trogocytosis than phagocytosis in the initial days after macrophages infiltrate the spheroid. The subsequent increase in phagocytosis is consistent with cancer cells dying due to trogocytosis, then their corpses being phagocytosed, or with trogocytosis enabling phagocytosis. Currently, there are no tools to specifically block trogocytosis without impacting phagocytosis as well, since the molecules involved in trogocytosis appear to also be required for phagocytosis. Future studies will need to test how trogocytosis impacts phagocytosis and antibody efficacy.

## Video legends

**Video 1: Her2 CAR macrophage phagocytoses a SKOV3 cell**

Video shows Her2 CAR GFP (green) macrophage phagocytosing a SKOV3 cell (mCh-CAAX). Images were acquired on a spinning disc confocal every 4 minutes. Scale bar is 20 μm. Stills from this movie are shown in Figure 1.

**Video 2: Her2 CAR macrophage trogocytoses a SKOV3 cell**

Video shows Her2 CAR GFP (green) macrophage trogocytosing a SKOV3 cell (mCh-CAAX). Images were acquired on a spinning disc confocal every minute. Scale bar is 20 μm. Stills from this movie are shown in Figure 1.

**Video 3: Her2 CAR macrophages infiltrate a SKOV3 spheroid**

Video moves through the Z planes of a SKOV3 (mCherry-CAAX; magenta) incubated with Her2 CAR GFP (green) macrophages. The magenta line highlights the boundary of the spheroid in each slice. Scale bar denotes 40 μm. Images of a live, unfixed sample were acquired on a spinning disc confocal 24 hours after the sample was encapsulated in matrigel with macrophages. The distance between Z slices is 10 μm and the total depth is 120 μm.

**Video 4: GFP-CAAX macrophages infiltrate a SKOV3 spheroid**

Video moves through the Z planes of a SKOV3 (mCherry-CAAX; magenta) incubated with GFP-CAAX (green) macrophages. The magenta line highlights the boundary of the spheroid in each slice. Scale bar denotes 40 μm. Images of a live, unfixed sample were acquired on a spinning disc confocal 24 hours after the sample was encapsulated in matrigel with macrophages. The distance between Z slices is 10 μm and the total depth is 120 μm.

**Video 5: Her2 CAR macrophage phagocytoses a SKOV3 cell in 3D spheroid**

Video shows Her2 CAR GFP (green) macrophage phagocytosing a SKOV3 cell (mCh-CAAX) in a 3D spheroid. Images were acquired on a spinning disc confocal every 6 minutes. Scale bar is 20 μm. Stills from this movie are shown in Figure S1.

**Video 6: Her2 CAR macrophage phagocytoses a dividing SKOV3 cell**

Video shows Her2 CAR GFP (green) macrophage phagocytosing a dividing SKOV3 cell (mCh-CAAX). Images were acquired on a spinning disc confocal every minute. Scale bar is 20 μm.

Stills from this movie are shown in Figure 7.

## Methods

### Experimental models

#### Cell lines

SKOV3 and L929 cells were obtained from the ATCC. HL-60 cells were obtained from the Denise Montell lab at the University of California, Santa Barbara. HCT116 cells were obtained from the Chris Richardson lab at the University of California, Santa Barbara. PANC-1 cells were obtained from the Angela Pitennis lab at the University of California, Santa Barbara. Lenti-X 293T cells were purchased from Takara. SKOV3 and HCT116 cells were cultured in McCoy’s 5A (Fisher 16-600-108), 10% FBS, 1% PSG. PANC-1, L929, and 293T cells were cultured in DMEM (Fisher 11965118), 10% FBS, 1% PSG. HL-60 cells were cultured in IMDM (Fisher 12440061). Raji cells were cultured in RPMI with 1% Glutimax (Fisher 72400120), 10% FBS, 1% PSG. Cells were routinely tested for mycoplasma. Cells were used for <20 passages.

#### Lentivirus production

pMD2.G (Gift from Didier Trono, VSV-G plasmid, Addgene plasmid 12259), pCMV-dR8.2 (Gift from Bob Weinberg, Addgene plasmid 8455) (Stewart et al., 2003), and our target construct was cloned into the pHR backbone. Transgene plasmids were transfected with lipofectamine LTX (Invitrogen 15338-100) into HEK293T cells to generate lentivirus. The media was harvested at 72hr, drawn through a 0.45um filter, and concentrated in LentiX (Takara Biosciences 631232).

#### Bone marrow derived macrophage cell culture

Male and Female C57BL/6 mice between the ages of 6 to 10 weeks were sacrificed by CO2 inhalation. Hips, femurs, tibia, and shoulders were dissected and bone marrow was isolated according to Weischenfelt and Porse (Weischenfeldt & Porse, 2008). Bone marrow cells were differentiated in complete RPMI supplemented with 20% L929-conditioned media for 7 days in a humidified incubator with 5% CO2 at 37C. Fresh BMDM media was applied every 2-3 days. Macrophage progenitors were infected with the lentiviruses Her2 CAR, GFP-CAAX, or mCh-CAAX on day 5 post-harvest. Differentiated BMDMs were used for experiments from 7-10 days. Macrophage differentiation was confirmed by CD11b and F4/80 staining with >80% double positive.

#### Spheroid production

Low-adhesion dishes were prepared by coating the wells of a tissue culture treated 96 well plate (Fisher Scientific 087722C) with 1.5% agarose. SKOV3 cells were plated at 8,000 cells per well in normal growth medium supplemented with 2.5% of reconstituted basement membrane (rBM), Matrigel (Corning CB-40230C). Plates were centrifuged at 1000 rpm for 10 min at 4C to aggregate a plane of cells for spheroid formation. Cells were incubated for 72 hours at 37C upon which compact spheroids could be observed.

### Methods details

#### Plasmids

Her2 CAR GFP was constructed in the pHR vector as follows: Signal peptide-(MQSGTHWRVLGLCLLSVGVWGQD) derived from CD3ε; Extracellular-anti-HER2 (4D5-8) scFv (Carter et al., 1992); stalk and TM-aa 138–206 CD8 (Uniprot Q96QR6_HUMAN); R (single arginine insertion from cloning process); cytoplasmic domain (aa 45–86) of the Fc γ-chain UniProtKB - P20491 (FCERG_MOUSE); linker- GSGS; Fluorophore: mGFP.

H2B-iRFP contains H2B (Uniprot H2B1B_HUMAN) inserted into the pHR vector; Fluorophore iRFP_710_.

CDH1-eGFP contains CDH1 (AddGene 133805 (Toda et al., 2018)) inserted into the puromycin resistant and doxycycline inducible pCW57 vector (AddGene 71782 (Barger et al., 2019)); Fluorophore eGFP.

The GFP-CAAX in the pHR vector (Morrissey et al., 2018) contains eGFP fused to a C-terminal CAAX targeting motif: KMSKDGKKKKKKSKTKCVIM

mCh-CAAX in the pHR vector (Morrissey et al., 2018) contains mCh fused to a C-terminal CAAX targeting motif: KMSKDGKKKKKKSKTKCVIM

The iRFP-CAAX in the pHR vector (AddGene 170464 (Harris et al., 2021)) contains an iRFP_670_ fused to a C-terminal CAAX targeting motif: KMSKDGKKKKKKSKTKCVIM

#### Flow cytometry measurement of trogocytosis and phagocytosis

For adherent solid tumor cell lines, 35,000 cells were plated in 1 well of a 12 well dish and incubated overnight prior to the experiment. For suspended targets, including solid or diffuse tumors, 75,000 cells were directly added to 1 well of a 12 well dish at the time of experimentation. These plating conditions both resulted in ∼75,000 cell targets at the time of the experiment. For antibody opsonization, purified mouse anti-human CD47 (Biolegend 323102, RRID:AB_756132) was spiked into the wells of cancer cells at a concentration of 5ug/mL. To achieve a ∼1:1 ratio of effector to target cell ratio, 75,000 Her2 CAR or GFP BMDMs were added to each well. BMDMs and cancer cells were co-incubated for 2 hrs, harvested in 0.25% trypsin EDTA (Fisher 25200072), neutralized, then washed in cold PBS. Samples were analyzed in an Attune NxT (Invitrogen). Further analysis was conducted in FlowJo. BMDMs were gated by GFP+ single cells, then internalization of cancer cells was distinguished for trogocytosis by mCh+/iRFP- and phagocytosis by mCh+/iRFP+ events.

#### TImelapse microscopy of macrophage activity in 2D monolayers

Unless otherwise specified, SKOV3 and macrophage co-incubations were performed with adherent SKOV3 and macrophages added in suspension. 10,000 SKOV3 were plated in 1 well of a 96 well glass bottom Matriplate (Brooks, MGB096-1-2-LG-L) and incubated overnight to establish adhesion. SKOV3 count approximately doubled overnight, yielding 20,000 cells. To achieve a ∼1:1 effector to target cell ratio, 20,000 Her2 CAR or GFP BMDMs were added to each well the following day. Co-incubations were imaged using spinning disc microscopy (40 × 0.95 NA Plan Apo air) for 8-10 hours with time intervals of 6 min or less. Internalized cancer cells were classified for trogocytosis or phagocytosis in ImageJ. Trogocytosis was classified as BMDMs that ingested fragments of the target cell membrane. Phagocytosis was classified as BMDMs that engulfed the target cell whole.

In figures 5A and S3, suspended SKOV3 cells were added to adherent macrophages at the start of the timelapse experiment at a 1:1 ratio of 20,000 macrophages:20,000 targets.

Mitotic mCh-CAAX SKOV3 were classified by the retraction of membrane to a round mother cell, formation of a cleavage furrow or the appearance of a daughter cell. Phagocytosis of a mitotic target was verified by any of the indicated hallmarks of mitosis.

#### Spheroid size measurements

After 72 hours on low adhesion plates, the resulting SKOV3 spheroids were transferred to a Matrigel dome of 50:50 rBM and NGM. BMDMs were added at 500,000 cells per dome to the liquid Matrigel. Domes were incubated at 37C for 30min to set the matrix before 500uL of NGM was added to the well. Whole spheroids were removed from the incubator and imaged daily for 10 days via spinning disc microscopy (10 × 0.95 NA Plan Apo air). Large images were captured in Nikon Elements by stitching together a grid of 4 images using the large image tool (15% overlap). The diameter of mCherry signal (mCherry-CAAX from SKOV3 cells and internalized mCherry in macrophages) was measured in both the X and Y dimension in ImageJ. The X and Y diameter was averaged to obtain the final spheroid diameter.

#### Quantification of trogocytosis and phagocytosis in spheroids

Spheroid macrophage co-incubations were imaged via spinning disc microscopy (40x 1.15 NA WI). BMDM activity was characterized in ImageJ by using mCherry-CAAX, H2B-iRFP SKOV3 cells to distinguish trogocytosis (mCherry+,iRFP-) and phagocytosis (mCherry+, iRFP+). 100um Z-stacks were captured at 2um steps. Only BMDMs that made contact with the spheroid were analyzed.

#### Macrophage invasion in spheroids

Using Imaris 10.1, 10x spheroid Z-stacks were cropped to fit the spheroid dimensions, reducing excess background signal. The SKOV3 fluorescent membrane signal was thresholded to generate a reconstructed spheroid surface. The reconstructed surface was overlaid to each slice of the Z-stack for ease of visualizing macrophages that had invaded into the spheroid.

#### Her2 expression measurement

35,000 mCh-CAAX H2B-iRFP SKOV3 were plated in 1 well of a 12 well dish and incubated overnight prior to the experiment. 75,000 Her2 CAR or GFP BMDMs were added to each well except single cell controls. BMDMs and cancer cells were co-incubated for 2 hrs, harvested in 0.25% trypsin EDTA, neutralized, then blocked in incubation buffer (PBS + 0.5% BSA) for 10 min. Cells were resuspended in incubation buffer for 30 min with either Brilliant Violet 421 isotype control (BioLegend 400157; RRID:AB_10897939) or Anti-CD340 BV421 (BioLegend 324420; RRID:AB_2563990). Samples were triple washed in PBS then analyzed in an Attune NxT. Further analysis was conducted in FlowJo to isolate the cancer cells and assess the median fluorescence intensity of each sample.

#### RGD peptide treatment

For RGD peptide treated cells, 20,000 SKOV3 cells were seeded in media supplemented with 50ug/mL of cilengitide trifluoroacetate salt (VWR International 88968-51-6). Media was aspirated and replaced with fresh media immediately before adding 20,000 macrophages. Trogocytosis and phagocytosis were measured by timelapse microscopy as described above.

#### Focal adhesion CRISPR knockout generation and validation

Focal adhesion knockouts were generated by digesting the guide scaffold of LentiCRISPR v2 (Addgene plasmid 52961) and replacing it with the top guides of ITGB1, ITGB5, ITGAV, and a non-targeting guide from the GeCKO v2 library (Table S1 (Sanjana et al., 2014). mCh-CAAX H2B-iRFP SKOV3 were infected with the lentivirus, and infected cells were selected with puromycin.

Integrin knockouts were validated for absence of surface expression via antibody staining. The SKOV3 integrin knockout cell lines were harvested in 0.25% trypsin EDTA, neutralized, washed in cold PBS, then blocked in incubation buffer for 10 min. Cells were resuspended in incubation buffer for 30 min with the following antibodies: FITC ITGB5 IgG1 kappa (Thermo 11-0497-41, RRID:AB_2043843), FITC isotype control IgG1 kappa (Thermo 11-4714-81, RRID:AB_470021), FITC ITAV/CD51 IgG2a kappa (BioLegend 327907, RRID:AB_940558), or FITC isotype control IgG2a kappa (BioLegend 400210, RRID:AB_326458). Samples were triple washed in PBS then analyzed in an Attune NxT. Analysis was conducted in FlowJo to assess the median fluorescence intensity of each sample.

#### E-cadherin overexpression

iRFP_670_-CAAX or H2B-iRFP_710_ Raji were infected with CDH1-eGFP and underwent puromycin selection to select for infected cells. 20,000 cells were plated into 1 well of a 12 well. To achieve the E-cadherin overexpression, cells were treated with doxycycline for 72 hr prior to the experiment. At the time of the experiment, control iRFP_670_-CAAX or H2B-iRFP_710_ Raji were added at 450,000 cells to 1 well of a 12 well, adjusted to the count of the E-cadherin overexpression cells. Cells were treated with purified mouse anti-human CD47 (Biolegend 323102, RRID:AB_756132) at a concentration of 5ug/mL. 450,000 BMDMs were seeded into each well and co-incubated for 2hrs. BMDMs and cancer cells were co-incubated for 2 hrs, harvested in 0.25% trypsin EDTA, neutralized, then washed in cold PBS. Samples were analyzed in an Attune NxT. Further analysis was conducted in FlowJo. BMDMs were gated by area/aspect ratio, mCh+ events. Within the mCh+ BMDM gate, phagocytosis was determined by the iRFP+ events from H2B-iRFP_710_ samples, whereas total activity was determined by the iRFP+ events from iRFP_670_-CAAX.

#### Cell cycle arrest

Cells were treated with 0.1uM paclitaxel (ThermoFisher P3456) or 1uM STLC (Sigma 164739-5G) for 20 hr prior to the experiment. To quantify cell cycle arrest, SKOV3 cells were infected with the FUCCI cell cycle reporter mCherry(G1)/BFP(S/G2) (Addgene plasmid 132429)(Sato et al., 2019). mCherry+ cells were considered to be in G1 and BFP+ cells were counted as S/G2/M. Timelapse imaging was used to measure phagocytosis as described above.

#### Phosphatidylserine staining

To measure phosphatidylserine, cells were stained with Pacific Blue Annexin (BioLegend 640918, RRID:AB_1279046). 35,000 SKOV3 cells were plated into 1 well of a 12 well and incubated overnight before treatment with a mitotic inhibitor for 20 hours. SKOV3 treated with 250uM Hydrogen peroxide for 20 hours served as a positive control for death. Cells were harvested in 0.25% trypsin EDTA, neutralized, washed in cold PBS, then blocked in incubation buffer for 10 min. Cells were resuspended in annexin binding buffer for 30 min with Pacific Blue Annexin. Samples were triple washed in PBS then analyzed in an Attune NxT.

#### Microscopy

Fluorescent images were acquired on a spinning disc confocal microscope (Nikon Ti2-E inverted microscope with a Yokogawa CSU-W1 spinning disk unit and a Hamamatsu Orca Fusion BT scMos camera). Objectives included 10 x 0.45 NA Plan Apo air, 40 × 0.95 NA Plan Apo air, 40 x 1.15 NA water immersion, and a 100 × 1.49 NA oil immersion. The microscope is equipped with a piezo Z drive. Temperature, CO2, and humidity were controlled via an OkoLabs stage top incubator. Image acquisition was controlled using Nikon Elements.

Brightfield images were acquired on an inverted ECHO Revolve with the 10 x 0.35 NA Plan Apo air objective. Image acquisition was controlled using ECHO software.

#### Statistical analysis

All statistical analyses were performed in Prism 10 (GraphPad) and presented as mean values and standard error of the mean. Figure legends indicate the statistical test performed and the number of biological replicates. In flow cytometry experiments, three technical replicates were averaged to calculate the biological replicate.

## Supporting information

Source data for figure 2a

## Acknowledgements

We thank the members of the Morrissey Lab for their constructive feedback and advice. We also thank Ryan Stowers for advice on the 3D culture system. This work was supported by NIGMS R35 GM146935 and University of California Cancer Research Coordinating Committee seed grant C23CR5592. M.M. is a Nadia’s Gift Foundation Innovator of the Damon Runyon Cancer Research Foundation (DRR-85-25). K.R. was supported by the UCSB Graduate Division Graduate Research Mentorship Program Fellowship and UCSB Graduate Division Research Accelerator Award. We acknowledge the use of the NRI-MCDB Microscopy Facility (Imaris) supported by the NIH: 1S10 ODO010610-01A1. Illustrations were prepared in BioRender.

