## Supplemental Figures for "Target cell adhesion limits macrophage phagocytosis and promotes trogocytosis"

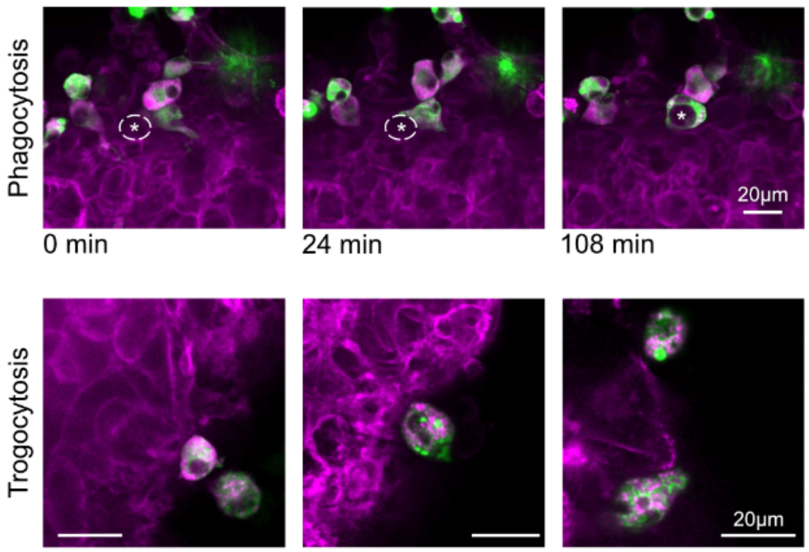


**Supplemental Figure 1 : Macrophage phagocytosis and trogocytosis in 3D Spheroids**

Top row shows a Her2 CAR GRP (green) macrophage phagocytosing a SKOV3 (mCherry-CAAX) cell in a 3D spheroid. The images are stills from Video 5. The dashed line and the asterisk denote the cell that will be phagocytosed. Scale bar is 20 um. Bottom row shows still images of Her2 CAR GFP (green) macrophages that trogocytosed, as evidenced by the small bites of SKOV3 (mCherry-CAAX) cells within the macrophage.

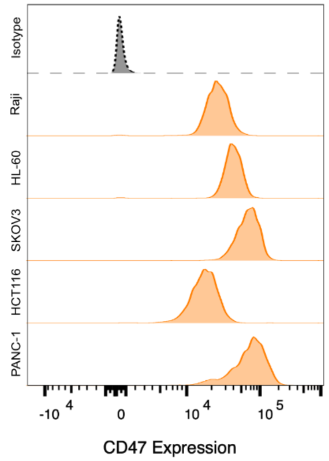


**Supplemental Figure 2 : CD47 expression on cancer cell lines**

The cancer cell lines used in Figure 4 were stained with a CD47 antibody and analyzed by flow cytometry. CD47 antibody binding was compared to an isotype control (grey).

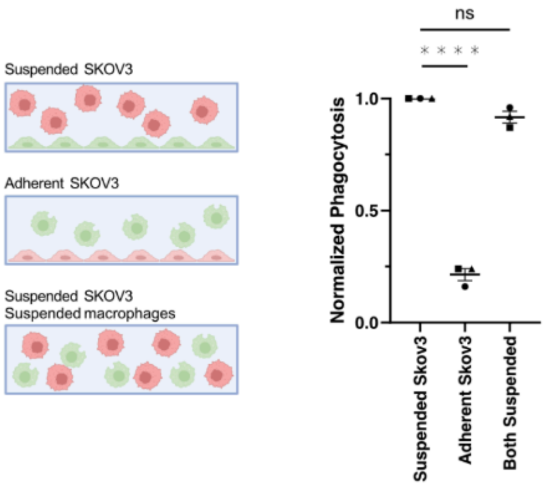


**Supplemental Figure 3: Adhesion of SKOV3 limits phagocytosis**

SKOV3 cells (red) labeled with mCherry-CAAX were incubated with Her2 CAR GFP macrophages (green) and phagocytosis was measured during a 10 hour timelapse. The number of phagocytic macrophages was normalized to the maximum phagocytosis in each replicate to control for batch-to-batch variability in BMDM appetite. Schematic shows the experimental set up with suspended SKOV3 cells and adherent macrophages, adherent SKOV3 cells and suspended macrophages, or with both cells in suspension at the start of the timelapse.

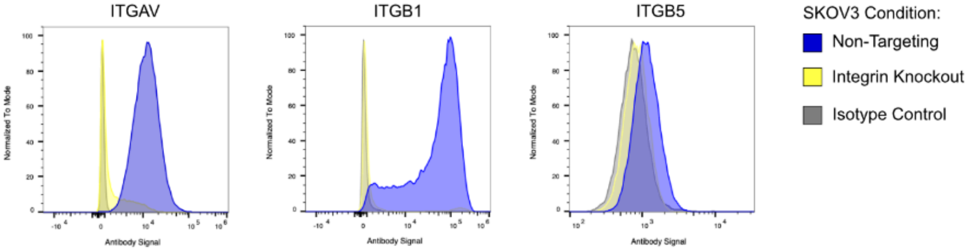


**Supplemental Figure 4: Validation of integrin knockout in SKOV3 cells**

SKOV3 cells were infected with a lentiviral construct encoding puromycin resistance, Cas9 and an sgRNA targeting ITGAV, ITGB1, ITGB5 or a non-targeting control. After puromycin selection, the polyclonal cell line was stained with an antibody targeting the relevant integrin subunit or an isotype control.

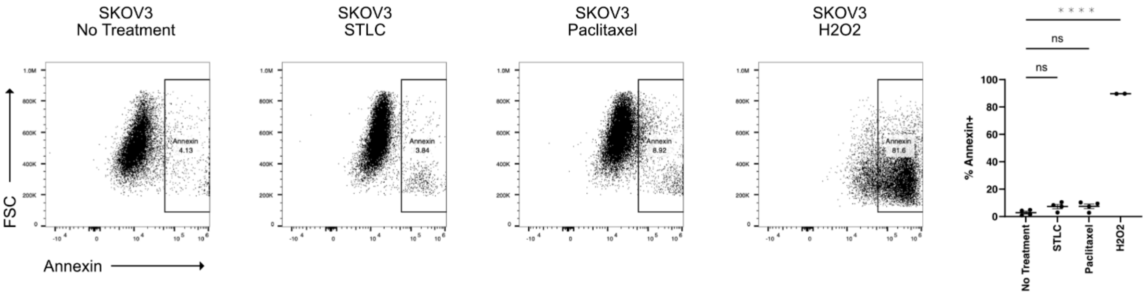


**Supplemental Figure 5: STLC and paclitaxel do not cause widespread phosphatidylserine exposure after 24 hours.**

SKOV3 cells were treated with STLC or paclitaxel for 20 hours, then stained with annexin to measure phosphatidylserine exposure, as an indicator of apoptosis. As a positive control, cells were treated with 250 μM H202 to induce apoptosis. Flow cytometry plots shows a representative experiment. Graph depicts the percent annexin positive cells with each dot representing an independent replicate.

**Supplemental Table 1: sgRNA Focal Adhesion Components**

Human GeckoV2 Library (Sanjana et al., 2014)

| Gene ID | Sequence |
|---|---|
| Non-targeting | ACGGAGGCTAAGCGTCGCAA |
| ITGB1 | ATCTCCAGCAAAGTGAAACC |
| ITGB5 | CCTCATCTCCTCCGCGAGTT |
| ITGAV | AGCATCTGTGAGGTCGAAAC |
